# SenPred: A single-cell RNA sequencing-based machine learning pipeline to classify senescent cells for the detection of an *in vivo* senescent cell burden

**DOI:** 10.1101/2023.10.23.563515

**Authors:** Bethany K. Hughes, Andrew Davis, Deborah Milligan, Ryan Wallis, Michael P. Philpott, Linda J. Wainwright, David A. Gunn, Cleo L. Bishop

## Abstract

Senescence classification is an acknowledged challenge within the field, as markers are cell-type and context dependent. Currently, multiple morphological and immunofluorescence markers are required for senescent cell identification. However, emerging scRNA-seq datasets have enabled increased understanding of the heterogeneity of senescence. Here we present SenPred, a machine-learning pipeline which can identify senescence based on single-cell transcriptomics. Using scRNA-seq of both 2D and 3D deeply senescent fibroblasts, the model predicts intra-experimental and inter-experimental fibroblast senescence to a high degree of accuracy (>99% true positives). We position this as a proof-of-concept study, with the goal of building a holistic model to detect multiple senescent subtypes. Importantly, utilising scRNA-seq datasets from deeply senescent fibroblasts grown in 3D refines our ML model leading to improved detection of senescent cells *in vivo.* This has allowed for detection of an *in vivo* senescent cell burden, which could have broader implications for the treatment of age-related morbidities.

## Introduction

Senescence is a fundamental cellular programme defined as a stable cell cycle arrest. The lack of universal senescence markers is a well-recognised challenge within the field. Hallmarks of senescence are both cell-type and trigger dependent, and no single marker alone is sufficient to confirm senescence in all contexts^1,2^. It is therefore necessary to assess multiple markers encompassing cellular morphology, effectors of cell cycle arrest, the senescence associated secretory phenotype (SASP), mitochondrial changes, chromatin remodelling, and lysosomal markers such as senescence associated beta galactosidase (SA-β-Gal)^1–3^. However, choosing appropriate markers for a particular experiment often relies on either literature precedent using the specific cell type and trigger, or lengthy optimisation and marker validation. Recent work has sought to address this by focusing on the development of accessible, stepwise, and standardised workflows for the classification and subsequent characterisation of senescence^2,4^. Kohli *et al.* proposed a two-step approach to senescence detection, firstly focusing on more general lysosomal changes (SA-β-Gal) and cell cycle arrest (lack of EdU incorporation, and immunocytochemistry for Ki67, p16 and p21), and secondly on subtype specific proinflammatory SASP changes^4^.

This seminal work has allowed for a more systematic and streamlined approach to senescence detection. The outcome of these approaches continues to depend upon the senescence context, meaning prior knowledge of the subtype is required to determine a senescent-cell state. Understanding the nuanced changes between senescence subtypes could allow for context-specific senescence detection and would enable unbiased classification of datasets where the senescent cell status is unknown. Furthermore, this understanding would remove the likelihood of detecting false positive and false negative data, which is crucial for progression within the senescence field. With the emergence of high-throughput screening methodologies, and ‘omics’ based sequencing technologies, our understanding of senescent cell subtypes is rapidly expanding. This necessitates the development of tools which can utilise this data to classify senescence in novel datasets.

Machine learning (ML) methodologies are widely used to classify distinct cellular states^5^. The data must have a state of ground-truth, so prior knowledge of the input class is required. The data can then be divided into training and testing datasets. The training dataset can be used to build the model, and the unseen testing dataset can be used to evaluate model performance based on a variety of metrics such as classification accuracy^6^. These ML models can then be applied to classify novel datasets. ML has been successfully applied to senescence classification in a small number of contexts (reviewed in Hughes, Wallis & Bishop 2023^5^). For example, Heckenbach *et al.* have developed their model ‘Xception’ using neural networks, which can accurately classify a cell as senescent based on its nuclear morphology^7^. Alongside morphology, RNA sequencing data has also been used to build predictive models based on mRNA expression levels of senescent versus proliferative cells^8–11^. Jochems *et al.* used an elastic net model to accurately classify therapy-induced senescent cancer cells grown in a 2D monolayer^11^. Importantly, the group stratified the cells into those which had undergone short- and long-term treatments and found that treatment time strongly influenced the transcriptome of the cells. When applying their model to *in vivo* patient derived xenografts treated with the senescence inducer SHP099, all xenografts were classified as ‘not senescent’. However, the group describe that the use of bulk RNA sequencing datasets could explain why they were potentially unable to detect senescence signatures *in vivo*.

Therefore, we asked if using single-cell transcriptomics alongside ML could improve senescent cell detection, particularly within heterogeneous *in vivo* contexts. Initially, we performed single-cell RNA sequencing of deeply senescent human dermal fibroblast (HDF) cells from a two-dimensional monolayer to build a ML model of senescence. We established that the model can accurately detect fibroblast senescence in independent 2D *in vitro* datasets, and in line with work by Jochems *et al.*, we demonstrate that temporal kinetics is an important consideration when building transcriptomic senescent cell models. Interestingly, we find that fibroblasts grown in a two-dimensional monolayer cannot generate an accurate model of *in vivo* senescence. Therefore, to more faithfully replicate conditions *in vivo,* we performed single cell RNA sequencing of cells grown in a three-dimensional matrix. We demonstrate that the 3D ML model enabled the detection of senescent fibroblasts *in vivo*. Here, we present SenPred, a proof-of-concept pipeline, which could facilitate the generation of a holistic ML model, encompassing a multitude of senescence contexts.

## Results

### Generating and evaluating machine learning models of replicative fibroblast senescence

In an effort to develop a ML model of senescence classification we cultured human dermal fibroblasts (HDFs) to replicative senescence (passage 39) and allowed them to reach a deeply senescent state over a further three weeks in culture (DS; P39+3). Single cell RNA sequencing (scRNA-seq) was carried out on Early Proliferative (EP) and Deeply Senescent (DS) HDFs cultured in two-dimensional monolayer (extensive model validation previously reported in Wallis *et al.*^5^) (Figure 1a). Using k-nearest neighbours (KNN), seven independent clusters were identified in the EP and DS HDF dataset (Figure 1b). Overlaying these clusters with the originating cell populations showed distinct clustering of EP and DS fibroblasts, highlighting the discrete mRNA profiles of the two populations, together with inherent heterogeneity within both the EP and DS fibroblast populations (Figure 1c). Given that the EP and DS states were transcriptionally distinct, we proceeded to build a ML model to distinguish the two conditions.

**Figure 1.**
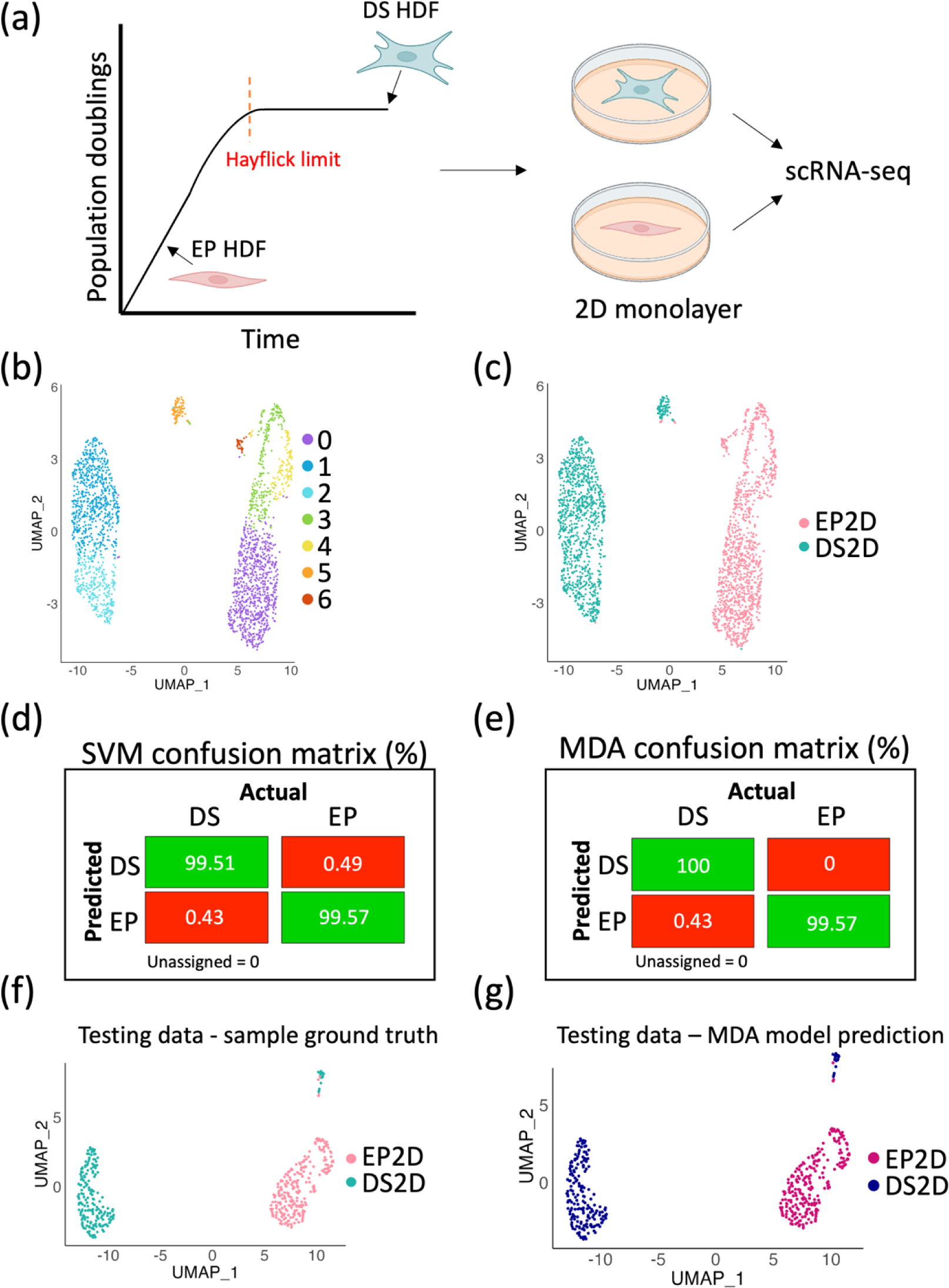
Machine learning models can accurately discriminate between Early Proliferative (EP) and Deeply Senescent (DS) human dermal fibroblasts. (a) Schematic illustration the position of Early Proliferative (EP) and Deeply Senescent (DS) Human Dermal Fibroblasts (HDFs) on the Hayflick curve. The samples were grown in a 2D mon olayer before single-cell RNA sequencing (scRNA-seq). (b) Uniform Manifold Approximation Projection (UMAP) of transcriptomic signatures of EP and DS fibroblasts. (c) UMAP in (b) overlayed with sample type. (d) Support Vector Machine (SVM) model confusion matrix generated from testing dataset, highlighting true positives and true negatives (green), and false positives and false negatives (red). (e) Multiple Discriminant Analysis (MDA) model confusion matrix generated from testing dataset, highlighting true positives and true negatives (green), and false positives and false negatives (red). (f) 20% testing dataset clustered into a UMAP and overlayed with sample type. (g) UMAP in (f) overlayed based on prediction using MDA model, with the ScPred package.

The scRNA-seq data was split into training (80%) and testing (20%) datasets, for model development and evaluation. Four ML models were explored, including Support Vector Machine (SVM; Figure 1d) and Multiple Discriminant Analysis (MDA; Figure 1e). The MDA confusion matrix highlighted a marginally higher percentage of true positive DS cells (0.49% higher than the SVM model). Both the SVM and MDA models have an ROC score of 1, and a sensitivity of 0.999, but the MDA had higher specificity than SVM (1 versus 0.99, respectively). KNN and Generalised Linear Model (GLM) were also tested but had marginally lower evaluation metrics (ROC: 0.998, Sens: 0.978, Spec: 0.992; and ROC: 0.999, Sens: 0.996, Spec: 0.994, respectively). For this reason, the MDA model was selected for subsequent work. Comparing the known classification of the testing dataset (Figure 1f), with the MDA model predictions (Figure 1g) shows a striking prediction accuracy of 99.79%. Although intra-experimental, this testing data is previously ‘unseen’, suggesting model overfitting is unlikely and instead the two classification outcomes are transcriptionally distinct. This work demonstrates that it is possible to build a MDA ML model of senescence classification, which can accurately predict fibroblast deep senescence in two-dimensional monolayer.

### Testing the models on an external dataset reveals importance of temporal senescence kinetics

To test the MDA ML model’s performance, we used an independent publicly available dataset from Chan *et al*. (GEO - #GSE175533), which included scRNA-seq data for increasing population doubling levels (PDLs) of WI-38 fibroblasts^13^. These fibroblasts ranged from PDL25 to PDL50, the latter of which Chan *et al*. described as being in an early senescent state. The transcriptional differences of the PDL50 senescent cells were apparent when the cells are clustered and plotted onto a UMAP (Figure 2a), with the majority of the PDL46 and PDL50 cells clustering away from the main bulk of the population. Applying our MDA ML model to the WI-38 fibroblast dataset shows an enrichment for DS predictions in the UMAP locations of the early senescent PDL50 cells (Figure 2b). However, only 19.19% of the PDL50 cells are predicted to be DS (Figure 2c). To further investigate this, the PDL50 cells were isolated and clustered independently, producing eight clusters (Figure 2d). Intriguingly, cluster five was largely enriched for DS predicted cells, suggesting that this subpopulation of cells were transcriptionally similar to the DS cells in the original HDF model.

**Figure 2.**
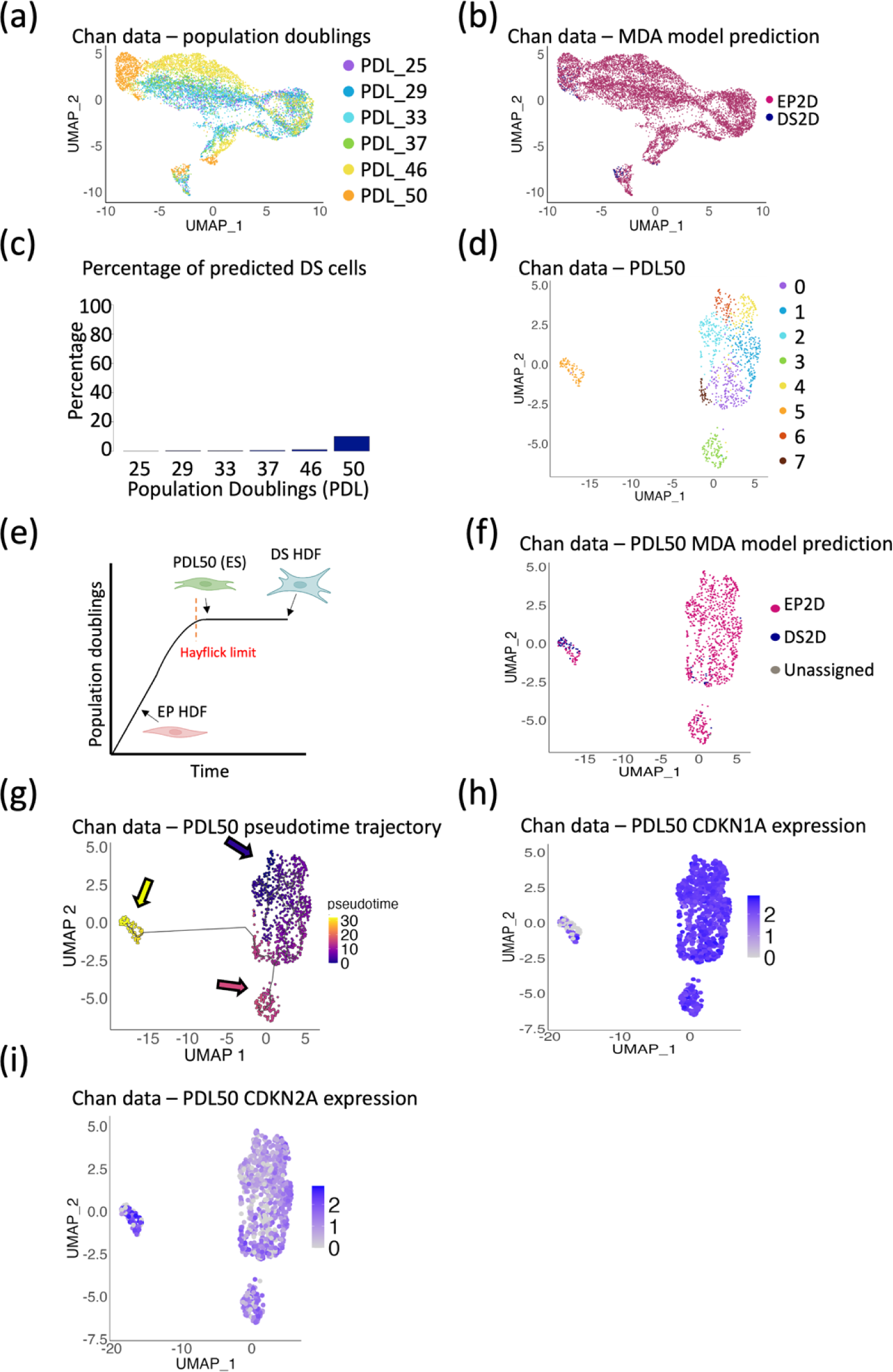
The machine learning model can detect deeply senescent but not early senescent fibroblasts. (a) UMAP of scRNA-seq data from fibroblasts from external dataset (Chan et al. ^13^) at different Population Doubling Levels (PDL). (b) UMAP in (a) overlayed based on prediction from the MDA model. (c) Barplot representing the p ercentage of cells which are predicted to be Deeply Senescent (DS) at each PDL, using the MDA model. (d) UMAP of the PDL50 cells from Chan et al. ^13^. (e) Schematic illustrating position on the Hayflick curve of Early Proliferative (EP) and Deeply Senescent (DS) Human Dermal Fibroblasts (HDFs), and PDL50 or ‘early senescent’ cells. (f) UMAP in (d) overlayed based on prediction from the MDA model. (g) Monocle3 trajectory plot of UMAP in (d), overlayed with predicted pseudotime trajectory. (h) PDL50 UMAP from (d), overlayed with expression of CDKN1A. (i) PDL50 UMAP from (d), overlayed with expression of CDKN2A.

We hypothesised that the low percentage of DS prediction in the PDL50 cells could be due to temporal differences in the two experiments. The DS cells used to build the MDA ML model had undergone three weeks of culture after reaching replicative senescence, to establish and deepen their phenotype. Conversely, the PDL50 WI-38 fibroblast cells from Chan *et al*. were sequenced immediately upon reaching their Hayflick limit (Figure 2e). To test this hypothesis, we applied Monocle3 trajectory analysis to the PDL50 cells^14^. Cluster five (the cluster enriched for cells which were predicted to be DS – Figure 2f), fell at the end of the predicted trajectory, suggesting that the model was detecting deeply senescent but not early senescent fibroblasts (Figure 2g) within the PDL50 population. Cells at PDL50 likely represent a heterogeneous population, where some cells have reached the Hayflick limit earlier and deepened their senescent phenotype. To confirm this, we examined the expression of known cell cycle arrest markers CDKN1A (p21) and CDKN2A (p16). Alcorta *et al.* reported an initial increase of p21 during the early stages of cell cycle arrest of human diploid fibroblasts, with a gradual increase of p16 expression as the senescence phenotype deepens^15^. Intriguingly, this was also apparent in the PDL50 cells, with those cells predicted to be DS having lower expression levels of CDKN1A (Figure 2h) and higher expression levels of CDKN2A mRNA (Figure 2i) than the remaining PDL50 population. This supports our hypothesis that the MDA ML model is able to detect deeply senescent but not early senescent fibroblasts.

To build a ML model which could encompass early replicative senescent fibroblasts (ES), the PDL50 cells from Chan *et al.*^13^ were incorporated into the model as an additional ES classifier. First, clustering the ES, EP, and DS cells revealed a more successive pattern, with the ES cells bridging the gap between the EP and DS cells (Figure 3a-b). This was confirmed using Monocle3 trajectory analysis (Figure 3c). This is perhaps unsurprising, as we have previously identified the ES cells to contain a heterogeneous mix with a small proportion of DS cells. The MDA confusion matrix highlights more than 90.2% true positives for all intra-experimental predictions, with the lowest percentage of true positives being for the ‘early senescent’ or ES cells (Figure 3d), which could again be explained by the heterogeneity of this population. Applying this model, the WI-38 cells at PDL25 through to PDL50 were able to be classified as either EP, ES, or DS (Figure 2a, Figure 3e). Unsurprisingly, the PDL50 cells had the highest predicted percentage of ES cells (66.6%). The percentage of predicted PDL50 cells then incrementally decreased with decreasing PDL number, with 15.7% of PDL25 cells being predicted to be ES (Figure 3f). This suggests that even at the earlier passages, the population of fibroblasts is heterogeneous, and that individual cells are on their own trajectories. Finally, for completeness, this ML predictive model for ES, EP, and DS cells was tested on the EP and DS HDF cells alone, identifying a small percentage (10.06%) of the EP cells as early senescent (Figure 3g). However, within the DS cells, only 0.39% were predicted to be ES, indicating that the DS cells are a more homogenous population.

**Figure 3.**
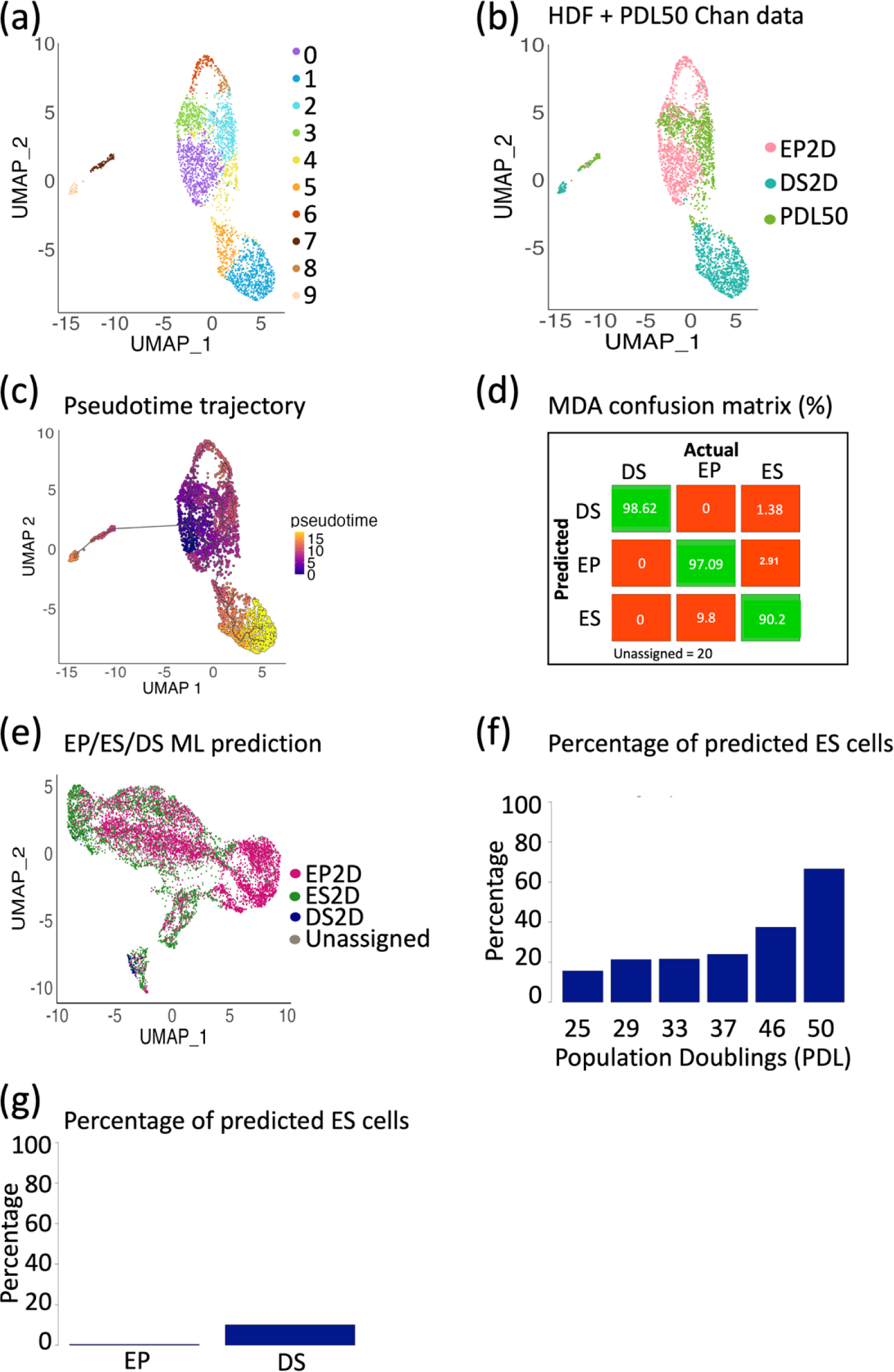
Building a machine learning model incorporating Early Senescent (ES) cells highlights high levels of heterogeneity throughout progressive population doublings. (a) UMAP of combined scRNA-seq data from EP and DS HDFs, and PDL50 WI −38 fibroblasts from Chan *et al.* ^13^. (b) UMAP in (a) overlayed with sample type. (c) Pseudotime trajectory of UMAP in (b). (d) Confusion matrix of MDA model of EP, ES, and DS cells, highlighting true positives and true negatives (green), and false positives and false negatives (red). (e) UMAP of all WI-38 PDLs from Chan *et al*.^13^ (Figure 2a), overlayed with prediction from EP/ES/DS model. (f) Barplot showing percentage of predicted ES cells at each PDL. (g) Barplot showing percentage of predicted ES cells within the EP and DS populations.

### Testing the models using *in vivo* data

To investigate whether the EP/ES/DS model can detect senescence *in vivo*, the models were tested on two independent publicly available whole skin scRNA-seq datasets from Tabib *et al.*^16^ (data kindly provided by corresponding author) and Sole-Boldo *et al*.^17^ (#GSE130973). Clustering the fibroblasts from Tabib *et al.* revealed five clusters (Figure 4a), which did not appear to stratify based on the age of the donors (Figure 4b), or by their predicted senescence state (Figure 4c). Importantly, the age of donor and senescent cell burden did not distinctly correlate. Even in the youngest donor fibroblasts, 63.1% of cells were predicted to be deeply senescent (Figure 4d).

**Figure 4.**
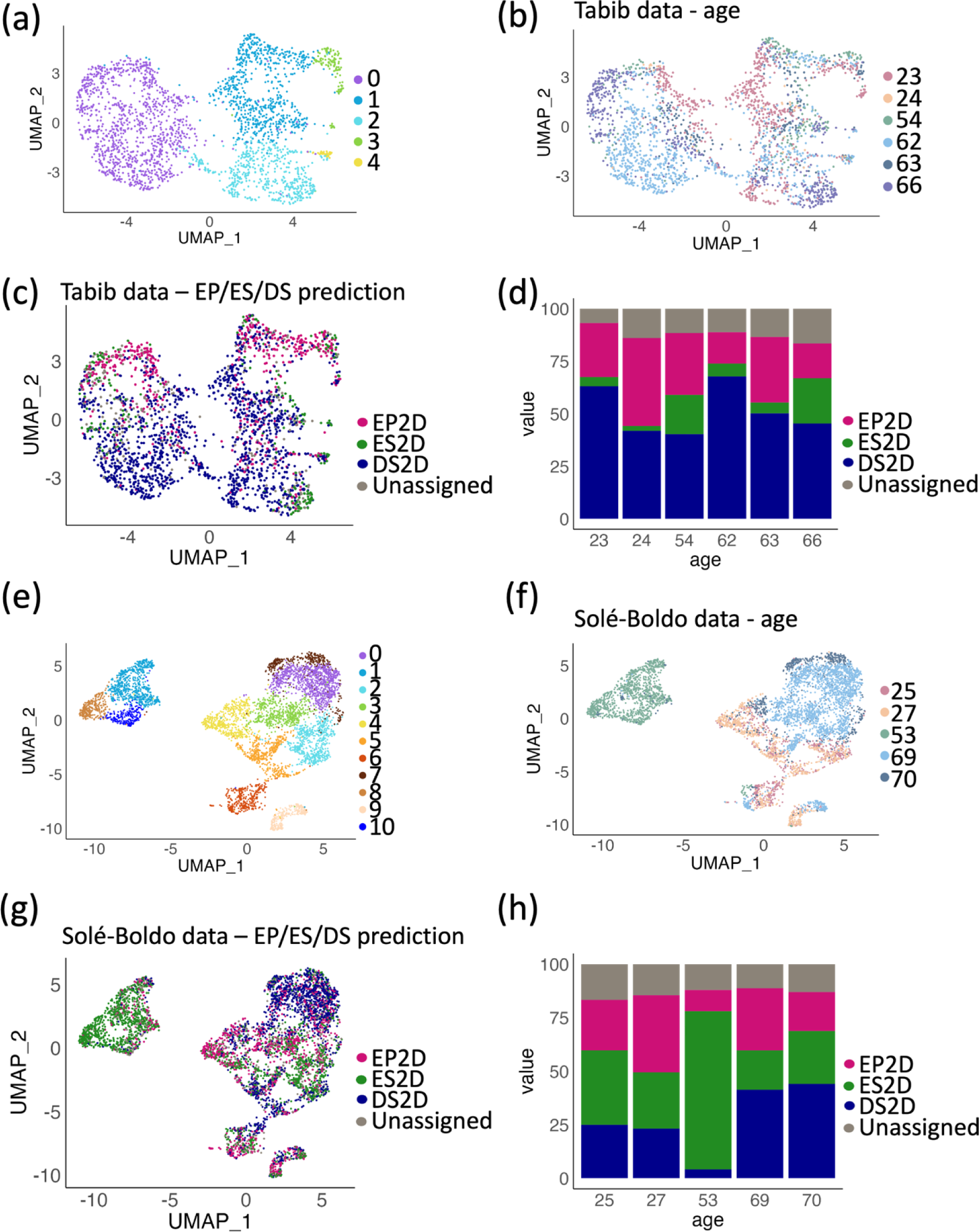
The machine learning models cannot accurately be applied to *in vivo* datasets. (a) UMAP of scRNA-seq of *in vivo* fibroblasts from Tabib *et al.* ^16^. (b) UMAP shown in (a), overlayed with donor age. (c) UMAP shown in (a), overlayed with MDA EP/ES/DS model prediction. (d) Barplot showing percentage model prediction of EP/ES/DS cells for each donor in (b). (e) UMAP of scRNA-seq of *in vivo* fibroblasts from Solé-Boldo *et al*.^17^. (f) UMAP shown in (e), overlayed with donor age. (g) UMAP shown in (e), overlayed with MDA EP/ES/DS model prediction. (h) Barplot showing percentage model prediction of EP/ES/DS cells for each donor in (f).

Analysis of the second whole skin scRNA-seq data from Sole-Boldo *et al*^17^ generated similar findings. Clustering these fibroblasts formed 11 distinct clusters (Figure 4e), and these did appear to have some age-associated influences (Figure 4f). Particularly, the 53-year-old donor clustered alone, which could suggest fibroblast abnormality for this individual, or perhaps issues with tissue isolation and processing. Intriguingly, this abnormality was also reflected in the percentage predictions of EP, ES, and DS cells, with the 53-year-old donor having a very high prediction of ES cells (Figure 4g-h). Again, although the proportion of predicted DS cells is slightly lower in the dataset from Sole-Boldo *et al.* compared with Tabib *et al.*, the combination of DS and ES predicted cells makes up a minimum of 49.44% of the fibroblasts in all donors. Evidence within the literature suggests that senescent fibroblasts within human skin *in vivo* make up approximately 10% of the fibroblast population^12^. The disparity between the model prediction and literature reports suggests that the model is not accurately able to identify senescent fibroblasts *in vivo*.

### Building *in vitro* 3D models improves senescent cell detection *in vivo*

In recent years, the limitations of culturing cells in a two-dimensional monolayer have been widely reported. Most importantly, cells cultured in a monolayer do not replicate the complexities and depth of the extracellular matrix *in vivo*, which can have wide implications on cell growth and signalling processes^18,19^. Consequently, three-dimensional organotypic models have been developed in an effort to recapitulate *in vivo* cell behaviour more accurately. For this reason, we built a 3D fibroblast matrigel-collagen dermal gel containing either EP or DS HDFs, which underwent 10X scRNA-seq (Figure 5a). Firstly, clustering of these cells revealed six distinct groups, and a clear separation of EP and DS cells (Figure 5b-c), highlighting the transcriptomic differences between EP and DS cells are maintained in three-dimensional culture. Next, a MDA ML model was built to classify the cells from the dermal gels as EP or DS, and intra-experimental testing revealed over 97.92% true positives (Figure 5d). Promisingly, applying this ML model to the dataset from Tabib *et al.*^16^ revealed a more physiological level of senescence prediction, with the maximum percentage of senescent fibroblasts being predicted in the 63-year-old (21.34%) (Figure 5e-f). It is important to highlight that the lack of ground-truth for the *in vivo* data means it was not possible to evaluate model accuracy *in vivo*., At present, this is likely to be an ongoing challenge when classifying *in vivo* data, due to the lack of a universal senescence marker *in vitro* or *in vivo*.

**Figure 5.**
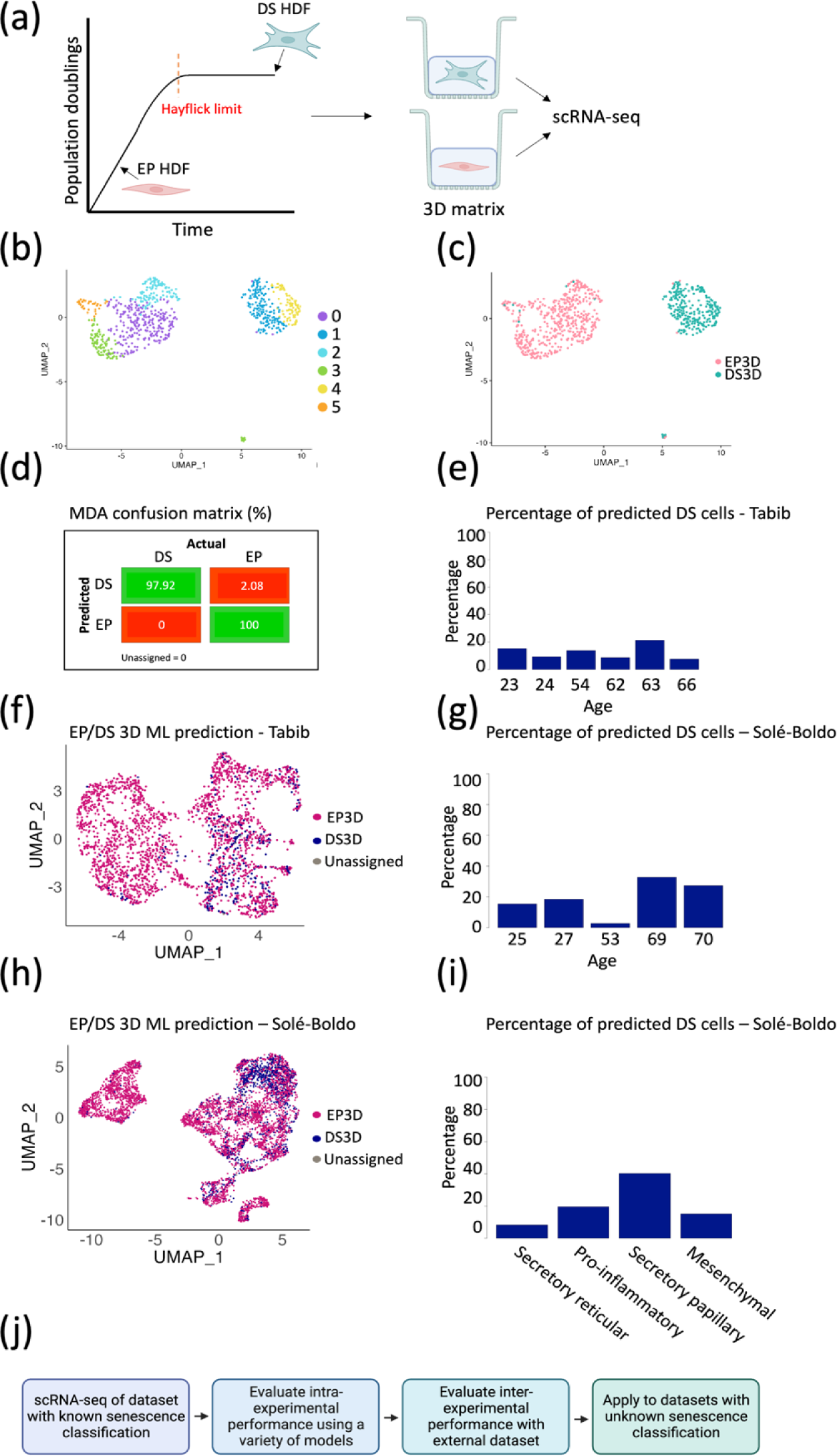
A ML model built from 3D dermal gels more accurately recapitulates senescence in vivo. (a) Schematic illustration the position of Early Proliferative (EP) and Deeply Senescent (DS) Human Dermal Fibroblasts (HDFs) on the Hayflick curve. The samples were grown in a 3D Matrigel-collagen matrix before single-cell RNA sequencing (scRNA-seq). (b) UMAP clustering of scRNA-seq of HDFs grown in a 3D matrigel matrix. (c) UMAP in (b), overlayed with sample type. (d) Confusion matrix of MDA model of EP and DS 3D cells, highlighting true positives and true negatives (green), and false positives and false negatives (red). (e) Barplot showing percentage of predicted DS cells for Tabib *et al.* ^16^ at each donor age. (f) UMAP in Figure 4 a, overlayed with 3D ML model prediction. (g) Barplot showing percentage of predicted DS cells for Solé-Boldo *et al*.^17^ at each donor age. (h) UMAP in Figure 4 e, overlayed with 3D ML model prediction. (i) Barplot showing percentage of predicted DS cells for Solé -Boldo *et al.* for each fibroblast subtype. (j) Flowchart showing the steps of the SenPred pipeline.

The three-dimensional ML model was also applied to the fibroblast data from Sole-Boldo *et al.*^17^. The predicted senescent cell burden in these individuals was higher than the individuals from the Tabib dataset (2.78-32.79% and 7.62-21.34%, respectively) (Figure 5g-h). The data from Sole-Boldo *et al.* indicated more of an age-related trend, with the older donors having a higher percentage DS prediction compared to the younger donors (27.36% for the 70-year-old donor compared to 15.47% for the 25-year-old donor) (Figure 5g). Again, the 53-year-old donor shows an abnormal profile, perhaps reflective of the clustering data in Figure 4f.

Mine *et al.* reported that papillary fibroblasts change with age compared to reticular fibroblast, including increasing secretion of keratinocyte growth factor, increased age-related contraction of collagen gels, and decreased population doublings and clonogenic capacity with age^20^, suggesting that papillary fibroblasts are likely to be the fibroblast subtype to undergo replicative senescence *in vivo*. Therefore, we investigated the proportion of senescent cells within the four fibroblast subtypes defined by Sole-Boldo *et al.*: secretory reticular, pro-inflammatory, secretory papillary, and mesenchymal. Promisingly, the secretory papillary fibroblasts from Sole-Boldo *et al.* had the highest percentage of senescence prediction (40.25%), providing increased confidence that the ML model is detecting senescent cells *in vivo* (Figure 5i).

In summary, the SenPred pipeline allows generation of an unbiased predictive model of senescence. Using scRNA-seq data from cells with a known senescence classification, ML models can be generated and evaluated both intra- and inter-experimentally (Figure 5j). These models can then be applied to classify senescence within unknown datasets. Importantly, using scRNA-seq data generated from cells grown in 3D allows detection of senescent cells *in vivo*.

## Discussion

The term ‘senescence’ encompasses a wide variety of cellular outcomes, which display diverse context-dependent states^1–3^. For this reason, there is no universal marker of senescence, and we believe that characterisation and classification of senescence should focus on identifying specific senescent cell subtypes. It is also important to consider the heterogeneity of these subtypes, and particularly the heterogeneity of *in vivo* aged tissue, where the minority of cells within the tissue are likely to be senescent^12^. The emergence of scRNA-seq datasets has allowed for greater understanding of this heterogeneity at the transcriptomic level. However, the lack of a single established senescence marker means identifying senescent cells within these datasets is arduous and requires prior knowledge of the specific senescent cell subtype. Recently, the SenNet Consortium was established (www.sennetconsortium.org), which aims to provide an atlas of senescence characterisation and mapping. With the growth of this database, it is important to have a tool which will allow rapid classification of senescence based on the characterisation atlas. Although AI and ML are emerging techniques for the classification of cellular subtypes, particularly within whole tissues or disease states, to date ML has not been used to classify senescence based on scRNA-seq data. Previously published work focussed on morphological classification, or using bulk RNA sequencing datasets which perhaps miss the nuanced nature of senescence^7–11^.

Within this work, we develop a ML model which can successfully classify deeply senescent fibroblast in both internal and external *in vitro* datasets. Applying the ML model based on scRNA-seq from *in vitro* data was unable to successfully determine senescence in fibroblasts from *in vivo* tissues. However, sequencing cells which were grown in a 3D environment more closely recapitulated reported senescent cell numbers *in vivo*, and also detected senescence in the expected fibroblast subtypes. Evaluation of model performance *in vivo* presents a clear limitation of this study, whereby the state of ‘ground truth’ is not known. The opportunity to use spatial transcriptomics alongside a classic senescence marker, although perhaps reductive based on the nuanced nature of senescence, may allow a more accurate evaluation of model performance *in vivo*. The advantages of multi-modal AI could be further explored, combining multiple ‘omics’ datasets to improve senescence prediction. The limited sample sizes of the *in vivo* datasets is a further limitation, with biological variability having a strong influence on the trend of senescence with age. With the emergence of larger datasets, this could improve the evaluation of the models and allow statistical power to detect directionality of senescence prediction. Physiological assessments of the donors would also strengthen the approach, as it would allow comparisons of biological versus chronological age.

We position this study as proof-of-concept, with the future aspiration to build a more holistic ML model in line with work by the SenNet consortium, whereby the model could detect multiple subtypes of senescence based on multiple triggers and cell types. This could then be applied to novel datasets and would allow rapid classification of the subtypes of senescence within tissues. A holistic senescence prediction model would have multiple clinical benefits, including predicting individual patient senescent cell burden and trajectory and the likelihood of senescence-associated age-related diseases. Alongside this, the model could be used to test senolytic activity by measuring clearance of senescent cells. This would support work by Smer-Barreto *et al.*, using ML to discover novel senolytics^21^. In conclusion, we believe ML is a valuable tool which can contribute towards the collective goal of the senescence field to characterise and classify senescence.

## Methods

### Cells and reagents

Primary human dermal fibroblasts (HDFs) were a kind donation from anonymous healthy patients under standard ethical practise, reference LREC No. 09/HO704/69. HDFs were cultured in DMEM with 4mM L-glutamine (41966-029, Life Technologies) supplemented with 10% foetal bovine serum (1001/500, Biosera), in the absence of antibiotics, and maintained at 37°C, 5% CO_2_, 95% humidity. Cells were seeded at 4,000 cells/cm^2^ for passage 1-19, 7,500 cells/cm^2^ for passage 20-25, and 10,000 cells/cm^2^ for passage 26 onwards. HDFs were serially passaged and were classified as Early Proliferative (EP) up to passage 13, and Deeply Senescent (DS) beyond passage 39+3. All cells were routinely tested for mycoplasma and were shown to be negative.

Dermal gels were generated by suspending HDFs in rat-tail collagen 1 (Corning) supplemented with Matrigel (Corning), FCS (1001/500, Biosera), 10X DMEM (11430-030, Invitrogen), and pH balanced with 4M NaOH in the absence of antibiotics. HDF-gel mix was added dropwise to transwells to give a final cell density of 10,000 HDFs per gel. Nine gels were constructed per condition. Dermal gels were maintained at 37°C, 5% CO_2_, 95% humidity.

### Single cell RNA sequencing

For 2D samples, cells were grown in 6-well plates at 7,000/cm^2^ for 72 hours. Cells were collected, and resuspended in HBSS (Thermofisher), and immediately sorted into single cell oil droplet suspension. To extract the cells from 3D dermal gels, gels were incubated with collagenase D (Sigma) at 37°C for 1 hr. Collagenase activity was subsequently inhibited by HBSS addition. Gels were strained and centrifuged to capture cell pellets, which were resuspended in HBSS before sorting into single cell oil droplet suspension.

Library generation and sequencing was performed using the v2 Chromium Single Cell 3’ Kit (10X genomics). Samples were sequenced using the Illumina NextSeq 500 High Output Run Sequencing platform using paired end sequencing and aligned to the human genome (GRCh37) using Cell Ranger (10x Genomics).

### Cell-level filtering and normalisation

All subsequent data analysis was performed using R (version 4.2.2) in RStudio (2022.07.2 Build 576), running under macOS Monterey 12.0.1. Matrix, gene, and barcode .gz data files were loaded and converted into a Seurat object^22^. Unless otherwise stated, default parameters from Seurat V4.3.0.1 were applied. To remove low quality cells, the data was filtered using the following metrics (Supplementary Figure S1):

- nCount_RNA > 1000
- nFeature_RNA > 400
- percent.mt <= 12

Data was normalised using LogNormalize, which normalises individual gene expression of a cell to total gene expression of this cell and multiplies by the default scale factor (10,000). This data is then log transformed.

### In vivo data cell-level filtering and normalisation

Cell level filtering and normalisation of datasets from Tabib *et al*. and Solé-Boldo *et al.* was performed using Seurat R package version 3.2.2 in R version 4.0.3. To remove low quality cells, the data was filtered using the following metrics:

- nCount_RNA > 500
- nFeature_RNA > 250
- nFeature_RNA <7500
- percent.mt <= 5
- log10genes_per_UMI <0.8

Data was normalised using ScTransform^23^, and the sctransform normalized counts for each sample were then integrated^24^, using the top 3000 most variable genes in the SelectIntegrationFeatures() function. Then, the FindIntegrationAnchors() function was applied with default parameters to identify the anchors across the different samples in each dataset. The anchors were then passed to the IntegrateData() function to create a single normalized and integrated Seurat object. All subsequent *in vivo* data analysis was performed in line with *in vitro* data analysis.

### Dimensionality reduction and differential gene expression analysis

The top 2000 variable genes were identified using the FindVariableFeatures function^24^, and these were subsequently scaled to ensure a mean expression of 0 and variance of 1 across all cells. K-nearest neighbour algorithms were used based on PCA distance, followed by the Louvain algorithm to determine clusters. The resolution of all cluster-based functions was determined systematically using the Clustree package^25^. These clusters were visualised on UMAP plots. To identify differentially expressed genes (DEGs) in the clusters, FindAllMarkers default Seurat function was implemented. Genes must be detected in >25% of cells in at least 1 cluster and have a logFC greater than 0.25.

### Machine learning senescence classifier

The ScPred package developed by the Powell group was used to build ML models to classify senescence (https://github.com/powellgenomicslab/scPred/)^26^. Unless otherwise stated, default parameters of ScPred V1.9.2 were used. Data was separated into training (80%) and testing (20%) datasets. ScPred ML models were then built on training data using all principal components, using a variety of methods including Support Vector Machines with Radial Basis Function Kernel (SVM), Mean Decrease in Accurary (MDA), K-nearest neighbours (KNN), and Generalised Linear Model (GLM). The models were evaluated based on their classification of the testing dataset through a variety of methods, including ROC, sensitivity, and specificity, and the best performing model (MDA) was taken forwards. For the MDA model, a fivefold cross-validation was used. These models could then be applied to classify senescence in alternative datasets.

### Trajectory analysis

Monocle3 V1.3.1 was used to analyse the trajectory of the PDL50 cells^14,27,28^. All default parameters were used unless otherwise stated. The earliest principal node was calculated using the function get_earliest_principal_node, where the node most surrounded by EP cells was selected. The cells were then ordered based on pseudotime.

### Package dependencies

All packages and versions used in this code are available at the following URL: https://github.com/bethk-h/SenPred_HDF. (Please note, the repository is currently private but will be made public upon paper publication).

### Data Availability

To be submitted to suitable database upon publication. Data from Solé-Boldo *et al.*^17^ and Chan *et al.*^13^ was accessed using the Gene Expression Omnibus (GEO #GSE130973 - Solé-Boldo, GEO #GSE175533 - Chan). The pre-processed dataset, including the raw UMI data matrix and associated metadata from Tabib et al., was acquired from the corresponding author, Robert Lafyatis in 2018.

### Code Availability

All code for this project is accessible via the following URL: https://github.com/bethk-h/SenPred_HDF. (Please note, the repository is currently private but will be made public upon paper publication). Full code will be shared with reviewers upon request.

---

**Figure S1.**
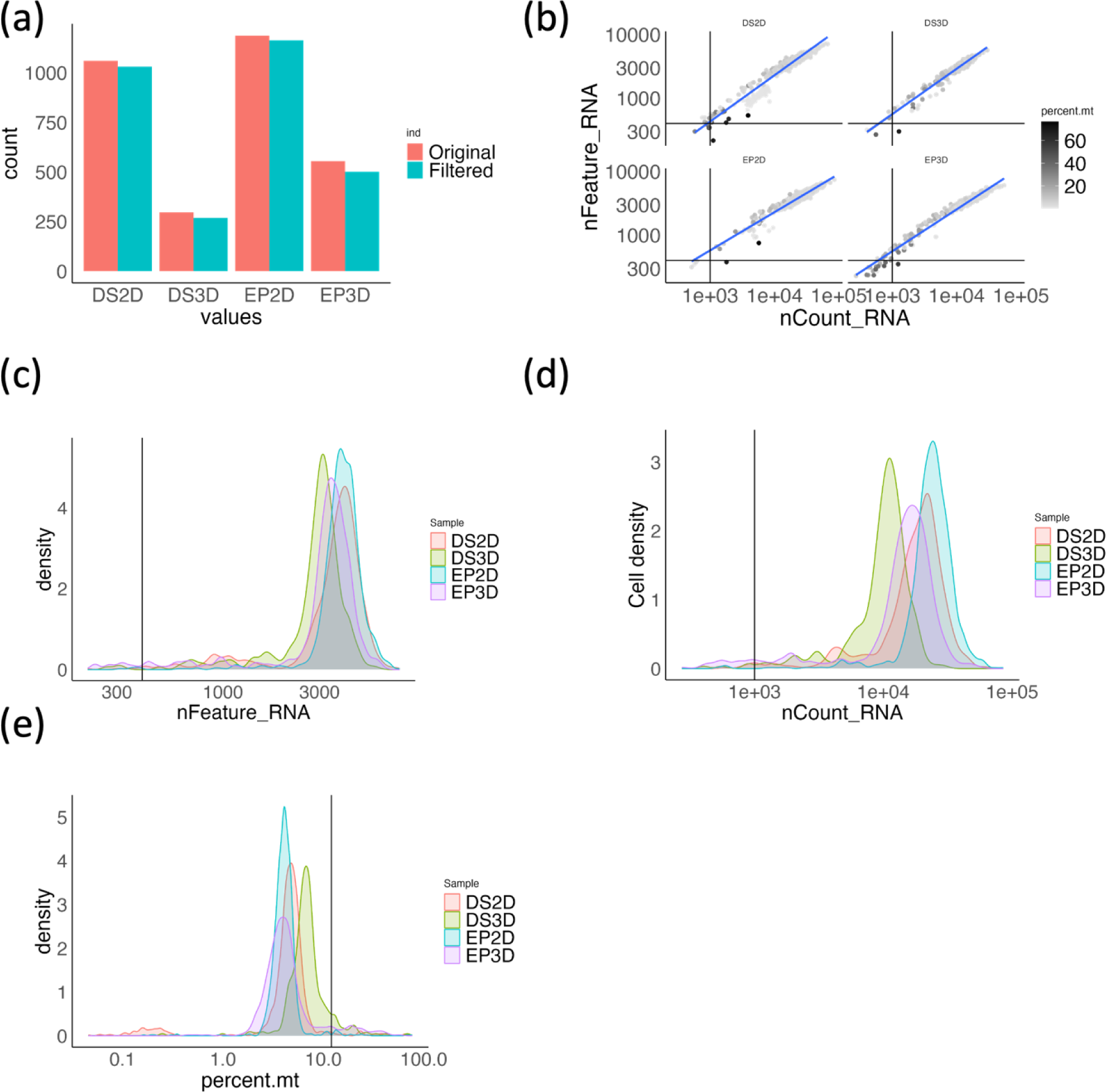
Quality control and filtering of scRNA-seq. (a) Number of cells sequenced before and after filtering. (b) Number of genes (n Feature_RNA) versus number of unique molecular identifiers (UMIs) (nCount_RNA), coloured by the percentage of mitochondri al RNA. (c) Density plot of nFeature_RNA for all samples. (d) Density plot of n Count_RNA for all samples. (e) Density plot of the percentage.mt for all samples. The black line represents chosen filtering cut offs in all cases.

